# Worth the weight: Sub-Pocket EXplorer (SubPEx), a weighted-ensemble method to enhance binding-pocket conformational sampling

**DOI:** 10.1101/2023.05.03.539330

**Authors:** Erich Hellemann, Jacob D. Durrant

## Abstract

Structure-based virtual screening (VS) is an effective method for identifying potential small-molecule ligands, but traditional VS approaches consider only a single binding-pocket conformation. Consequently, they struggle to identify ligands that bind to alternate conformations. Ensemble docking helps address this issue by incorporating multiple conformations into the docking process, but it depends on methods that can thoroughly explore pocket flexibility. We here introduce Sub-Pocket EXplorer (SubPEx), an approach that uses weighted ensemble (WE) path sampling to accelerate binding-pocket sampling. As proof of principle, we apply SubPEx to three proteins relevant to drug discovery: heat shock protein 90, influenza neuraminidase, and yeast hexokinase 2. SubPEx is available free of charge without registration under the terms of the open-source MIT license: http://durrantlab.com/subpex/

## 2. Introduction

Drug-discovery research and development costs range from $161 million to a staggering $4.54 billion per drug^1,2^. To improve efficiency, researchers leverage computational methods in most steps of the drug-discovery pipeline. Structure-based virtual screening (VS) is a common technique that can alleviate the high costs associated with early-stage hit identification. Given a virtual compound library, VS involves first docking each compound into a protein binding pocket to predict its pose (i.e., 3D geometry of binding). Second, each pose is assigned a score that (hopefully) correlates with affinity. Researchers select the most promising candidate ligands for further study.

Traditional VS methods dock flexible compounds into a static (rigid) binding pocket associated with a single protein-receptor structure. But in reality, binding pockets are flexible, continuously sampling multiple geometries, any of which may be amenable to small-molecule binding^3^. Researchers who consider only one pocket conformation may identify some ligands, but they risk discarding ligands that bind other valid conformations.

To address the shortcomings of rigid-receptor docking, many modern VS projects dock candidate ligands into multiple protein-pocket conformations^3-14^, an approach called multiple-conformation VS^3,15-21^ or ensemble docking. Structures obtained via experimental methods such as X-ray crystallography, NMR, and cryo-electron microscopy can provide useful conformations for ensemble docking^22^, but most drug targets are associated with only a limited number of experimental structures (if any), and those structures often capture very similar (i.e., redundant) pocket conformations.

Despite known limitations^23,24^, brute-force molecular dynamics (MD) simulations are a rich source of protein conformations that can effectively supplement experimental structures^17,19-21^, providing a more complete view of pocket flexibility^25-28^. In brief, MD simulations capture atomic movements by approximating the dynamics of molecular systems using classical (Newtonian) forces. Based on the positions and interactions between atoms, MD engines solve the equations of motion to iteratively update atomic positions over time, thus providing insight into the system’s behavior.

Many drug-discovery-relevant pocket dynamics occur on timescales accessible to standard (brute-force) simulations^16,17,19,21,25-28^. Such simulations sometimes reveal unanticipated side pockets (i.e., contiguous extensions of a primary pocket) that can bind novel chemical moieties^29^. But if considerable energy barriers separate druggable conformational states, standard MD may take too long to thoroughly sample the entire conformational landscape. Many conformational changes remain computationally inaccessible^30-32^.

In this work, we present <u>Su</u>b-<u>P</u>ocket <u>EX</u>plorer (SubPEx), a tool that uses weighted ensemble (WE) path sampling, as implemented in the Weighted Ensemble Simulation Toolkit with Parallelization and Analysis (WESTPA)^33^ framework, to accelerate protein-pocket sampling. SubPEx uses WE to efficiently direct sampling towards ever more distinct binding-pocket shapes, allowing users to enhance pocket sampling^3,20,21,28,34^ *in silico*, even without a known ligand. As proof of principle, we apply SubPEx to three proteins relevant to drug discovery: heat shock protein 90, influenza neuraminidase, and yeast hexokinase II. To encourage adoption, we release SubPEx under the permissive MIT open-source license. Users can obtain a copy free of charge without registration from http://durrantlab.com/subpex/.

## 3. Methods

### 3.1 Preparing proteins for simulation

We prepared three systems for simulation: the ATP binding domain of heat shock protein 90 alpha (HSP90; PDB 5J2V^35^), N1 neuraminidase (NA; PDB 2HU4 and 2HTY^36^ for closed and open conformations, respectively), and *Saccharomyces cerevisiae* hexokinase II (*Sc*Hxk2p; PDB 1IG8^37^). In the case of the 2HU4 N1 neuraminidase structure (see Supporting Information), we removed the bound ligand (oseltamivir).

After downloading the crystal structures from the Protein Data Bank^38^, we added hydrogen atoms to each protein using the PDB2PQR^39^ webserver (pH 7.0), which uses the PROPKA algorithm to optimize the hydrogen-bond network^40^. We used LEaP, part of the AmberTools18 package^41^, to add a water box that extended 10 Å beyond the protein in every direction. We also added Na+ or Cl-ions sufficient to neutralize the charge of the protein and then additional ions to approximate a 150 mM NaCl solution.

We simulated the systems with either the NAMD^42^ or Amber^41^ MD engines (Table 1). Regardless of the MD engine used, we parameterized all systems using the Amber ff14SB^43^ and TIP3P^44^ force fields for protein and water, respectively.

**Table 1.** Technical details for brute-force and SubPEx simulations. We used the minimal adaptive binning scheme^45^ for all SubPEx simulations listed here. See Table S1 for a description of addition simulations.

| Brute-force and SubPEX |  | SubPEX Only |  |  |
| --- | --- | --- | --- | --- |
| System | MD Engine | Progress Coordinate | # Bins | Walkers/Bin |
| HSP90 (5J2V)<sup>35</sup> | NAMD 2.14 | pRMSD | 19 | 3 |
| HSP90 (5J2V)<sup>35</sup> | NAMD 2.14 | cRMSD | 19 | 3 |
| NA (2HTY:A)<sup>36</sup> | Amber20 | cRMSD | 19 | 3 |
| Hxk2 (1IG8)<sup>37</sup> | NAMD 2.13 | cRMSD | 19 | 3 |

To resolve steric clashes, we first applied four rounds of minimization (5,000 steps each). In the first minimization step, we allowed hydrogen atoms to be free; in the second, hydrogen atoms and water molecules; in the third, hydrogen atoms, water molecules, and protein side chains; and in the fourth, all atoms. We followed each minimization with brief equilibration simulations. For the systems using NAMD, we used five serial equilibration simulations run in the NPT ensemble (250 ns each), gradually relaxing backbone restraints with each simulation (1.00, 0.75, 0.50, 0.25, and 0.00 kcal/mol/Å^2^, respectively). We used a 1 fs timestep in the first step and a 2 fs timestep in subsequent steps. For the systems using Amber, we used a three-step equilibration. We first simulated in the NVT ensemble for 10,000 steps, with a 2 fs timestep and 1 kcal/mol/Å^2^ backbone restraints; then in the NPT ensemble for 1 ns, with a 2 fs timestep and 1 kcal/mol/Å^2^ backbone restraints; and finally in the NPT ensemble for 1 ns, 2 fs timestep without any restraints.

### 3.2 Molecular dynamics and weighted ensemble simulations

For each system, we used the same MD engine to run subsequent brute-force molecular dynamics (MD) simulations. All simulations were run in the NPT ensemble, using a 2 fs timestep, the Monte Carlo barostat (pressure of 1.01325 bar, 1 atm), and the Langevin thermostat with a collision frequency of 5.0 ps^-1^ and a target temperature of 310K. For each system, we performed three brute-force production runs (500 ns, 250 ns, and 250 ns; 1 μs total).

To accelerate pocket conformational sampling, we performed SubPEx simulations of each system using the same respective MD engine, NPT ensemble, 2 fs timestep, Monte Carlo barostat, and Langevin thermostat. For all SubPEx simulations, we used a τ of 20 ps. We also used the minimal adaptive binning scheme^45^ for all SubPEx simulations but one (see Table S1). SubPEx accelerates molecular dynamics in a user-defined region (e.g., pocket) in 3D space. For each system, we defined an appropriate region by selecting residues that line a known binding pocket and visually verifying that the center of geometry of those residues fell within the confines of the pocket itself. Further details specific to each WE simulation (*e*.*g*., number of bins, walkers per bin) are given in Tables 1 and S1.

### 3.3 Simulation analyses

We calculated progress coordinates using in-house scripts. To load/manipulate all molecular data and to perform PCA analysis, we used the MDAnalysis python package (1.0.0)^46,47^. To cluster the MD trajectories, we used Amber20’s CPPTRAJ^48^ (hierarchical agglomerative clustering with average linkage^49^).

## 4. Results and discussion

### 4.1 Weighted-ensemble path sampling to capture rare events

SubPEx leverages the weighted ensemble (WE) path sampling method^50,51^ as implemented in the WESTPA 1.0 framework^33^ to enable conformational transitions over larger energy barriers. In brief, one first defines a uni- or multi-dimensional progress coordinate (i.e., measure of progress towards a target state), which is divided into discrete bins. Multiple stochastic, starting-bin simulations (“walkers”) then run in parallel for a predefined interval (τ), each carrying a statistical weight. At the end of each interval, one checks the simulations’ progress along the coordinate. To encourage even sampling, simulations that enter under-sampled bins are replicated with new seeds (kinetic energies) to improve sampling in that bin. Excess walkers that remain in oversampled bins are merged. The probabilities of the involved walkers are divided or added in the replicating and merging, respectively. WE thus accelerates rare-event sampling by encouraging even sampling along a predetermined progress coordinate. It focuses computational resources on under-sampled regions of conformational space^33^ without adding artefactual bias to the underlying free-energy profile.

### 4.2 SubPEx workflow

Running a SubPEx simulation involves several steps. First, the user must prepare a protein system that is fully parameterized, minimized, and equilibrated (i.e., ready for production MD simulation). The user also defines the protein pocket or region that will be subject to enhanced sampling. SubPEx measures the shape of the input pocket (described in greater detail below), which serves as an initial reference. It then launches multiple fixed-length, unbiased MD-simulation segments (walkers) from the initial conformation, as the WE approach requires.

At fixed time points (), SubPEx assesses the walkers’ progress along a carefully devised progress coordinate that captures the extent of pocket-shape dissimilarity relative to the initial measurement. Walkers that have made similar progress along the coordinate are grouped into progress-coordinate bins. SubPEx continuously replicates and merges walkers to encourage even sampling of all bins.

These steps continue iteratively until pocket-shape space is adequately sampled. Once the SubPEx run completes, the user clusters the sampled pockets to extract representative conformations (e.g., for use in ensemble virtual screening).

### 4.3 Methods for assessing pocket dissimilarity

SubPEx provides several methods for assessing pocket dissimilarity, as required to calculate its progress coordinate. We here describe two such methods; the Supporting Information includes additional descriptions.

#### 4.3.1 Pocket-heavy-atoms RMSD

SubPEx allows users to assess pocket dissimilarity by calculating the root-mean-square distance (RMSD) between the heavy atoms of the walker pockets and the reference pocket (pocket-heavy-atoms RMSD, or pRMSD). SubPEx identifies all pocket heavy atoms by considering every reference-structure residue within a user-defined distance of a given pocket-central point. It then calculates the pRMSD,

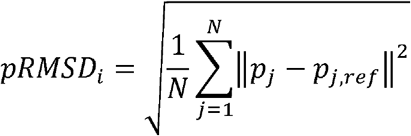

where *pRMSD*_*i*_ is the pRMSD value of walker *i, N* is the number of pocket heavy atoms, *p*_*j*_ is the position (in Euclidean space) of atom *j, p*_*j,ref*_ is the position of the same atom in the reference pocket, and ||*p*_*j*_ − *p*_*j,ref*_|| is the distance between those two atoms.

#### 4.3.2 cRMSD

In our tests, we found that progress coordinates that focus exclusively on the binding pocket sometimes sample only limited conformations. Simulations may only reach certain pocket conformations if the larger protein structure also undergoes at least subtle rearrangement. To focus (but not hyperfocus) computational resources on the pocket while allowing (and even encouraging) whole-protein conformational shifts, we developed the composite RMSD (cRMSD) progress coordinate:

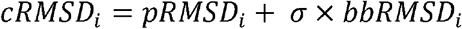

where *cRMSD*_*i*_ is the composite coordinate of walker *i*; *pRMSD*_*i*_ is the pocket-heavy-atom RMSD of walker *i*, defined above; *bbRMSD*_*i*_ is the similarly calculated RMSD of all protein backbone atoms; and σ is a proportionality constant. In our tests, we found that setting σ equal to the percentage of backbone atoms not in the pocket divided by two allowed SubPEx to effectively focus on pocket sampling while still permitting limited whole-protein backbone dynamics. The cRMSD coordinate is the SubPEx default.

### 4.4 Example of use: ATP-binding domain of HSP90

To demonstrate SubPEx utility, we performed SubPEx and brute-force MD simulations of the ATP binding domain of human heat shock protein 90 (HSP90). We chose HSP90 as a test protein because of its small size, excellent representation in the Protein Data Bank (260 structures with 100% sequence identity as of July 15^th^, 2022), and relevance to cancer therapy^52,53^. HSP90 is a molecular chaperone that plays a central role in many cellular processes, including cell-cycle control and survival. It is one of the most abundant proteins in the cytosol and helps maintain cellular homeostasis. HSP90 overexpression contributes to tumorigenesis^52-55^, so it is a well-studied chemotherapy drug target.

#### 4.4.1 pRMSD progress coordinate fails to adequately capture pocket flexibility

As an initial test, we began with an equilibrated *apo* HSP90 (PDB 5J2V^35^) structure and performed a SubPEx simulation using the one-dimensional pRMSD progress coordinate. The pRMSD SubPEx simulation ran for 50 generations (49.62 ns of cumulative simulation time and a maximal trajectory length of 1 ns). See Figure S2B for the pRMSD progress-coordinate values per WE generation.

To compare the pRMSD SubPEx simulations to brute-force simulations of the same system, we also ran three brute-force simulations of HSP90 (250 ns, 250 ns, and 500 ns, totaling 1 μs). We separately considered the first 50 ns of each brute-force simulation (to roughly match the cumulative simulation time of the SubPEx simulation) as well as the entire concatenated trajectory (1 μs) for reference. To assess the extent of pocket and backbone sampling, we calculated the pRMSD and bbRMSD of the simulated frames. We note that pRMSD and bbRMSD can also be used as SubPEx progress coordinates (see also the Supporting Information), but in this context, they serve as independent metrics (auxiliary data) to judge the extent of conformational sampling.

This analysis shows that the pRMSD SubPEx simulation sampled fewer pocket (and backbone) conformations than the similar-duration brute-force simulations (Figure 1**Error! Reference source not found**.). Counterintuitively, by focusing sampling on the pocket without simultaneously encouraging backbone flexibility, pRMSD SubPEx may not allow for the large-scale conformational changes required for some pocket rearrangements.

**Figure 1.**
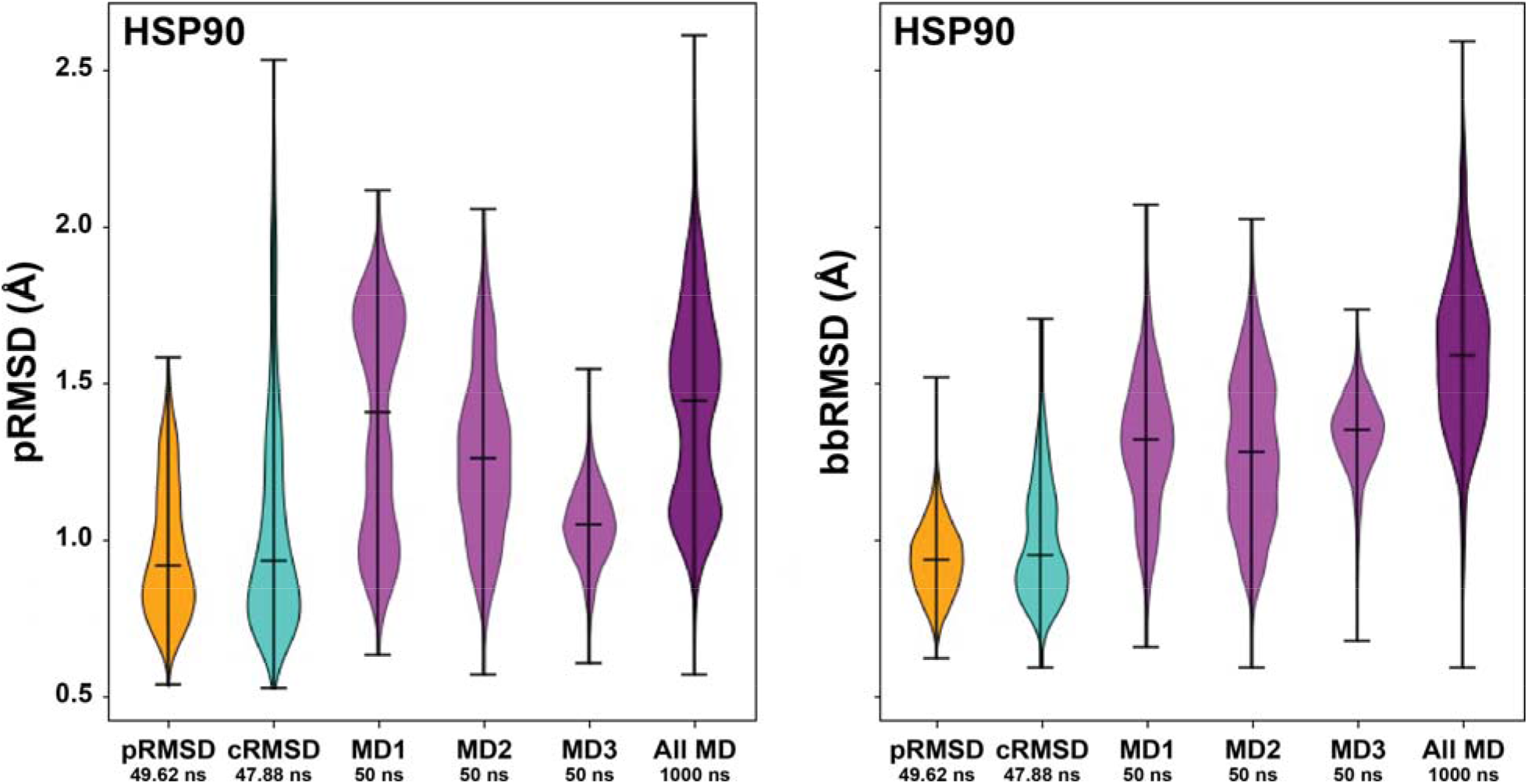
Violin plots comparing SubPEx and brute-force HSP90 simulations. The distributions of pRMSD values, indicative of pocket sampling, are shown on the left. The distributions of bbRMSD values, indicative of backbone sampling, are shown on the right. SubPEx sampling using the pRMSD and cRMSD progress coordinates is shown in orange and turquoise, respectively. Sampling by three brute-force MD simulations (first 50 ns) is shown in light purple. Sampling by a longer brute-force simulation (the three independent simulations, concatenated; 1 μs total) is shown in dark purple.

#### 4.4.2 cRMSD progress coordinate effectively enhances pocket flexibility

We next tested a progress coordinate that encourages some backbone flexibility while still focusing on the pocket: the one-dimensional cRMSD coordinate (a weighted linear combination of pRMSD and bbRMSD that favors pRMSD). We ran a cRMSD SubPEx simulation for 50 generations (47.88 ns of cumulative simulation time and a maximal trajectory length of 1 ns). See Figure S2C for the cRMSD values per WE generation.

The cRMSD SubPEx simulation sampled pockets with a far wider range of pRMSD values than the similar-duration brute-force simulations (**Error! Reference source not found**.Figure 1A), suggesting substantially enhanced pocket sampling. Remarkably, the cRMSD SubPEx simulation even sampled some pocket conformations with pRMSD values comparable to those of the much longer brute-force simulations (1 μs, concatenated). These measurements suggest cRMSD SubPEx dramatically enhances the number of identified pocket conformational states over the brute-force approach.

An analysis of the whole-protein bbRMSD provides insight into the source of this improvement (Figure 1B). The cRMSD SubPEx simulation does not sample the same range of bbRMSD values as the brute-force simulations, as expected given that SubPEx by design focuses conformational sampling on the pocket. But it does sample a wider range of bbRMSD values than the pRMSD SubPEx simulation, suggesting it is not as hyper-focused on the pocket at the expense of adequate backbone sampling. The cRMSD coordinate also performed better than other one-dimensional progress coordinates we tested on the HSP90 system (see Supporting Information).

### 4.5 SubPEx better samples physically relevant conformations

Having demonstrated that the cRMSD progress coordinate can effectively enhance pocket conformational sampling, we next verified that the HSP90 cRMSD SubPEx simulation better samples physically relevant conformations. We extended our previous 50-generation HSP90 cRMSD SubPEx simulation for an additional 50 generations (now 102.6 ns of cumulative simulation time vs. the 47.88 ns of the initial SubPEx run). We then performed principal component analysis (PCA^56^) on the pocket heavy atoms sampled by this extended cRMSD SubPEx simulation, three brute-force simulations each trimmed to the same length as the SubPEx simulation (102.6 ns), and a collection of 75 HSP90 crystal structures extracted from the Protein Data Bank. We concatenated all these conformations before calculating principal components that served as a common orthogonal basis set onto which the same conformations were then separately projected.

**Error! Reference source not found**.Figure 2 shows the cRMSD SubPEx simulation and the three brute force MD simulations projected onto the first and second principal components. In each panel, the crystal-structure projections are marked as red dots. We found that the SubPEx simulations sample the principal component space more thoroughly than the brute-force simulations (∼47% coverage of the PC space vs. ∼43%, 25%, and 11% coverage, respectively).

**Figure 2.**
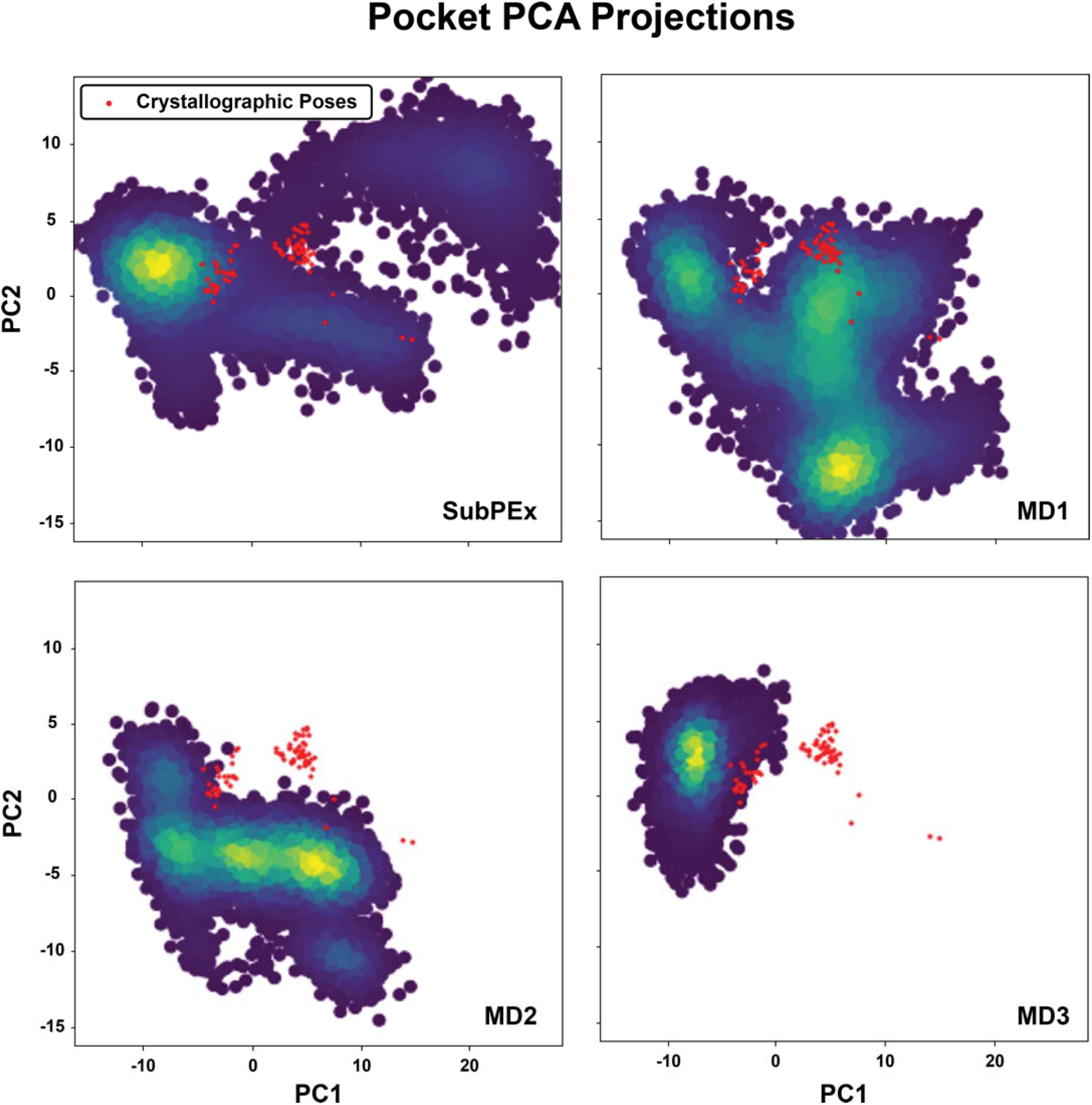
Principal component analysis of the cRMSD SubPEx and three brute-force HSP90 simulations (102.6 ns each), considering only pocket heavy atoms. All simulations were projected onto the same PC space, derived from the conformations of the SubPEx and brute-force simulations, as well as the selected crystallographic structures.

We noticed that the crystal structures projected onto this PCA space clustered mainly into two groups, with only a few outliers. Visual inspection revealed that the pocket-lining residues N105-A111 are primarily responsible for these two clusters. These residues are part of a larger stretch (D102-G114) that can form a helix (helix 3) or a loop, depending on the bound ligand^57^. As examples, consider the 4EFT^58^ and 4YKR structures, which capture D102-G114 in helical and loop conformations, respectively.

To assess how well the simulations captured the helical and loop conformations, we computed the pocket-heavy-atom RMSD between each simulation frame and the 4EFT and 4YKR structures, which served as crystallographic reference conformations. Although the SubPEx and brute-force simulations came within 2.3 Å of the helical conformation, neither captured that conformation exactly (e.g., < 1 Å). However, SubPEx more effectively sampleed regions both similar to and distant from the crystallographic references (Figure S3).

### 4.6 Simulation clustering to identify representative conformations

We next explored how best to extract representative conformations from SubPEx simulations for later use in ensemble docking. The traditional approach involves first striding a simulation (i.e., keeping only periodic, regularly spaced frames), clustering those strided conformations, and constructing an ensemble comprised of one conformation from each cluster (e.g., the centroids). Striding reduces the number of conformations that the clustering algorithm must consider, making it practical to calculate a pair-wise distance matrix that would otherwise be too computationally expensive. Striding is effective because standard MD simulations are linearly correlated in time. The frame sampled at timestep *t* + 1 is nearly identical to the one sampled at timestep *t* and so can be reasonably discarded as redundant. However, SubPEx-sampled conformations are not linearly correlated because they involve many branching simulations run in parallel. Striding is thus inappropriate.

To accelerate SubPEx clustering without striding, we used a two-tiered approach. We first used average-linkage hierarchical agglomerative clustering, as implemented in CPPTRAJ^48^, to generate a pocket-heavy-atom representative ensemble for each SubPEx generation. To avoid bias, the ensemble sizes per generation scale with the number of walkers. (1) The ensemble corresponding to the smallest generation contains *N* clusters, where *N* equals the number of walkers or three, whichever is larger. (2) The ensemble corresponding to the largest generation has twenty-five clusters. (3) Ensembles corresponding to intermediately sized generations scale proportionally. We next merge these many per-generation ensembles and again cluster the merged set. Calculating distance matrices per generation is much faster than considering all SubPEx-frames at once (Figure 3). Furthermore, this two-tiered approach is compatible with any clustering algorithm.

**Figure 3.**
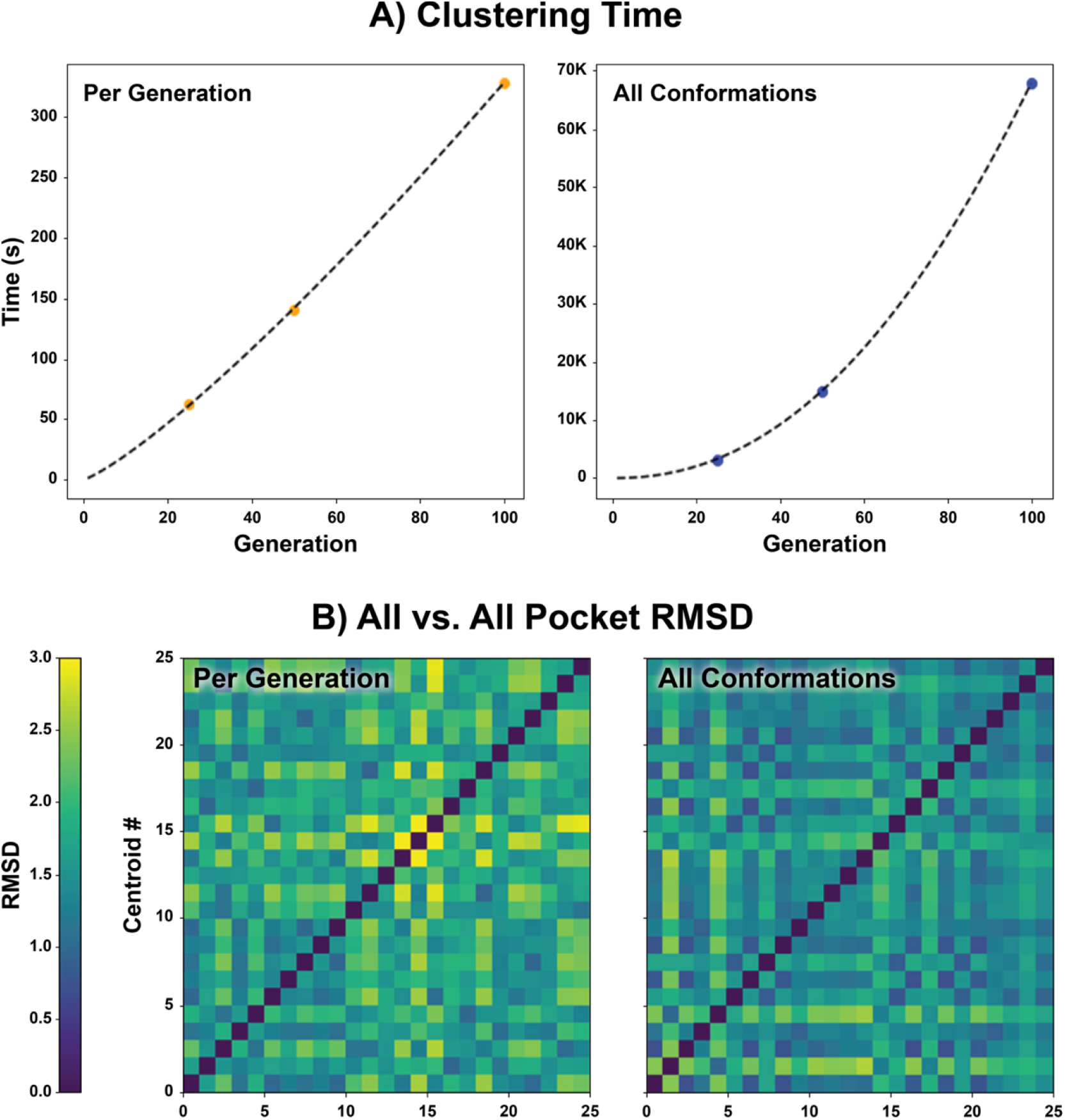
Comparison of per-generation and all-frame clustering (102.6 ns). (A) The time required to cluster per generation vs. using all frames. (B) All vs. all pocket RMSD of the centroids obtained from clustering. Results for per-generation and every-frame clustering are shown on the left and right, respectively.

To further confirm the enhanced pocket sampling of the cRMSD SubPEx simulation, we clustered the extended simulation (102.6 ns, 100 generations) using our per-generation approach. We separately clustered the first 102.6 ns of the third brute-force MD production run (same cumulative time) using traditional clustering but without any striding to ensure a fair comparison. Overlaying the centroids of these clusters provides a visual demonstration of the enhanced pocket diversity that SubPEx captures (Figure 4).

**Figure 4.**
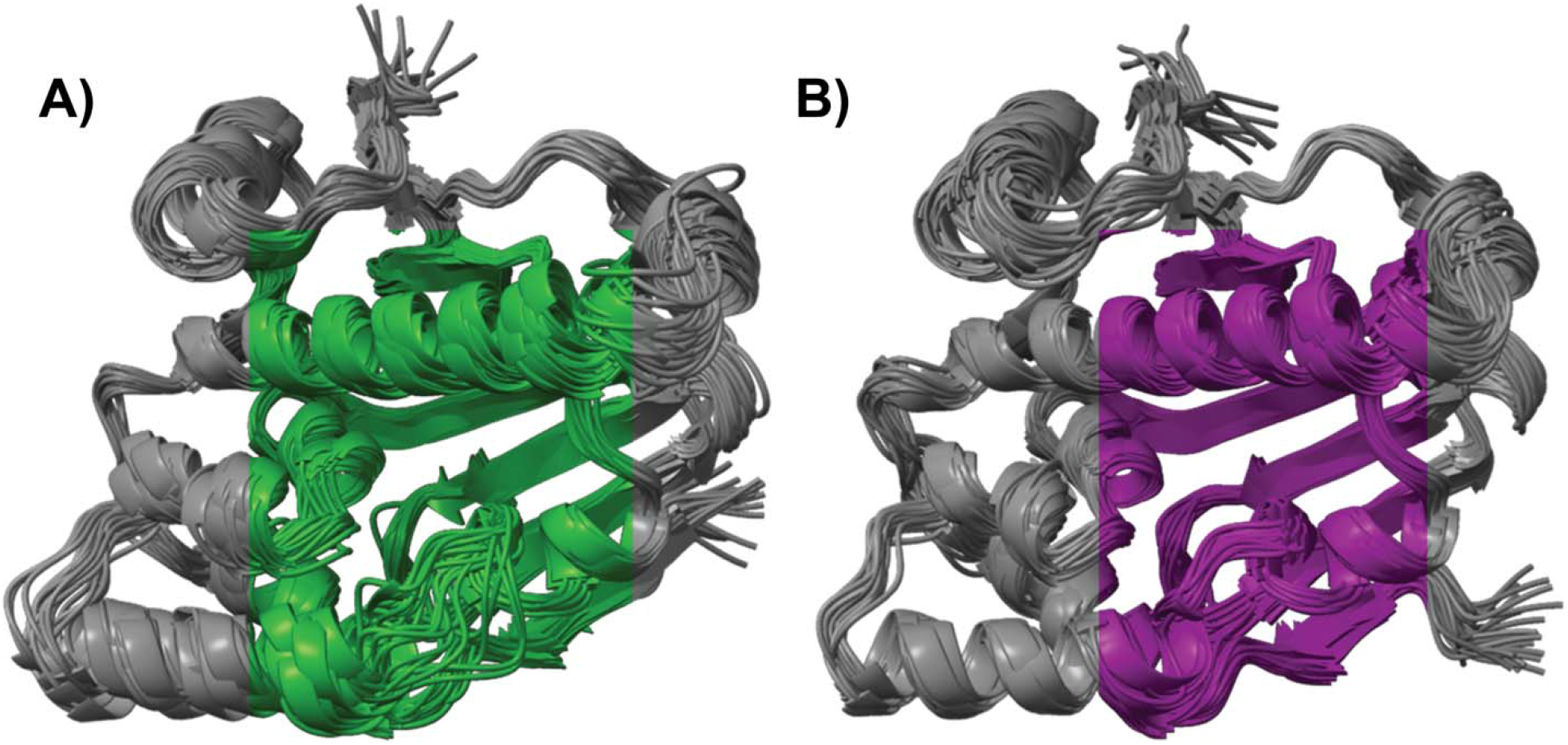
Superposition of HSP90 structures obtained from clustering SubPEx and brute-force simulations (same cumulative simulation time of 102.6 ns). (A) Structures sampled by the cRMSD SubPEx simulation. (B) Structures sampled by the brute-force MD simulations.

### 4.7 Example of use: influenza neuraminidase

To further demonstrate SubPEx utility, we performed SubPEx and brute-force MD simulations of neuraminidase, a protein with critical roles in the influenza infection cycle. Though most people recover from seasonal influenza after a couple of days, the virus still kills between 290,000 to 650,000 each year^59^. Additionally, influenza is responsible for occasional devastating pandemics (e.g., the 1918 Spanish Flu, which killed an estimated 50 million people^60^).

Following viral replication, influenza virions remain bound to the host cell by sialic-acid linkages that connect the viral hemagglutinin to the host-cell surface. Neuraminidase (NA) mediates the release of these virions by cleaving the sialic-acid linkages^61-63^. NA proteins have nine serotypes (N1 to N9), which can be divided into two groups according to their sequence. A main structural difference between the groups is the presence or absence of an extra cavity in the active site^64,65^. This cavity, which others have actively exploited for drug development^66^, is formed when the D147-H/R150 salt bridge breaks, increasing 150-loop flexibility^67^.

#### 4.7.1 cRMSD progress coordinate again effectively enhances pocket flexibility

To assess how well SubPEx improves sampling of the flexible 150-cavity, we performed three brute-force simulations (totaling 1 μs) of an NA system with an open-150 cavity (PDB 2HTY:A^36^). We also performed SubPEx simulations using the cRMSD progress coordinate, starting from the same open-150-cavity NA conformation (PDB 2HTY:A). SubPEx sampled pocket conformations with higher pRMSD values than the brute-force simulations while maintaining thorough (even) sampling over lower-pRMSD pocket conformations (Figure 5**Error! Reference source not found**., top row). The SubPEx simulations also sampled lower bbRMSD values than the brute-force simulations, showing that SubPEx still focused on enhancing pocket rather than whole-protein dynamics. Incorporating limited backbone flexibility into the progress coordinate, even if only modestly, again had an outsized impact on pocket sampling. See the Supporting Information for additional details regarding NA simulations that started from the closed-150-cavity conformation.

**Figure 5.**
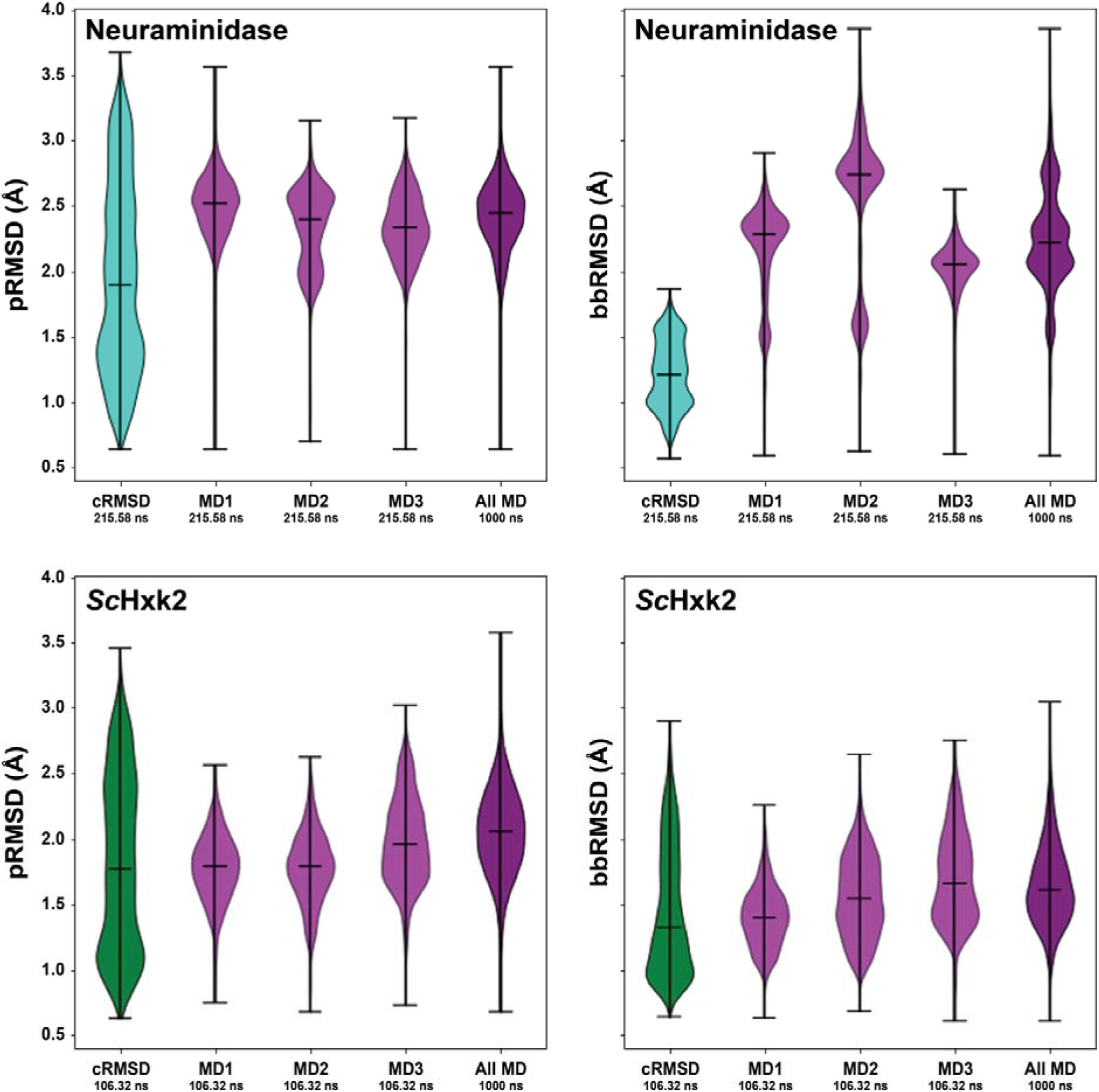
Violin plots comparing NA and *S*cHxk2 SubPEx and brute-force simulations. The distributions of pRMSD values, indicative of pocket sampling, are shown on the left. The distributions of bbRMSD values, indicative of backbone sampling, are shown on the right. **Top row**: **NA simulations**, starting from the open conformation. cRMSD SubPEx sampling is shown in turquoise. Sampling by three brute-force MD simulations (first 215.58 ns) is shown in light purple. Sampling by a longer brute-force simulation (the three independent simulations, concatenated; 1 μs total) is shown in dark purple. **Bottom row**: ***Sc*Hxk2 simulations**. cRMSD SubPEx sampling is shown in green. Sampling by three brute-force MD simulations (first 106.32 ns) is shown in light purple. Sampling by a longer brute-force simulation (the three independent simulations, concatenated; 1 μs total) is shown in dark purple.

### 4.8 Example of use: hexokinase II (Hxk2)

We next used *Saccharomyces cerevisiae* Hxk2 (*Sc*Hxk2p) to evaluate SubPEx on a system that undergoes large ligand-induced domain rearrangements. Hexokinases phosphorylate glucose at its sixth position^68,69^, allowing glucose to advance to downstream metabolic pathways. *Sc*Hxk2 contains a large and a small domain. In the unbound open state, the *Sc*Hxk2 enzymatic cleft is solvent exposed^70,71^. Glucose binding induces a relative rotation of the two subdomains^72,73^ that encloses or “embraces” the glucose molecule^74,75^. ATP binding then induces additional structural changes to complete the transition to the closed state^76^.

We performed both SubPEx and brute-force simulations of *Sc*Hxk2 starting from the open (glucose-absent) conformation (PDB 1IG8^37^; Figure 5**Error! Reference source not found**., bottom row). For the SubPEx simulations, we again used the cRMSD coordinate and ran for 100 generations (106.32 ns of cumulative simulation time and a maximal trajectory length of 2 ns).

We next used our two-tiered per-generation approach to cluster the SubPEx ensemble, and traditional clustering with striding (10,632 evenly spaced frames) to cluster the brute-force simulation. This clustering confirms that the SubPEx simulations sampled a wider range of conformations than the brute-force simulations and that those conformations are biologically relevant (Figure 6 and Figure S6). Though both simulations started from the open form (PDB 1IG8, in red), the SubPEx simulations also sampled conformations similar to the closed form (*Kl*Hxk1, PDB 3O8M^77^, in blue). These results suggest SubPEx can capture diverse pocket conformations even when those conformations require substantial conformational shifts.

**Figure 6.**
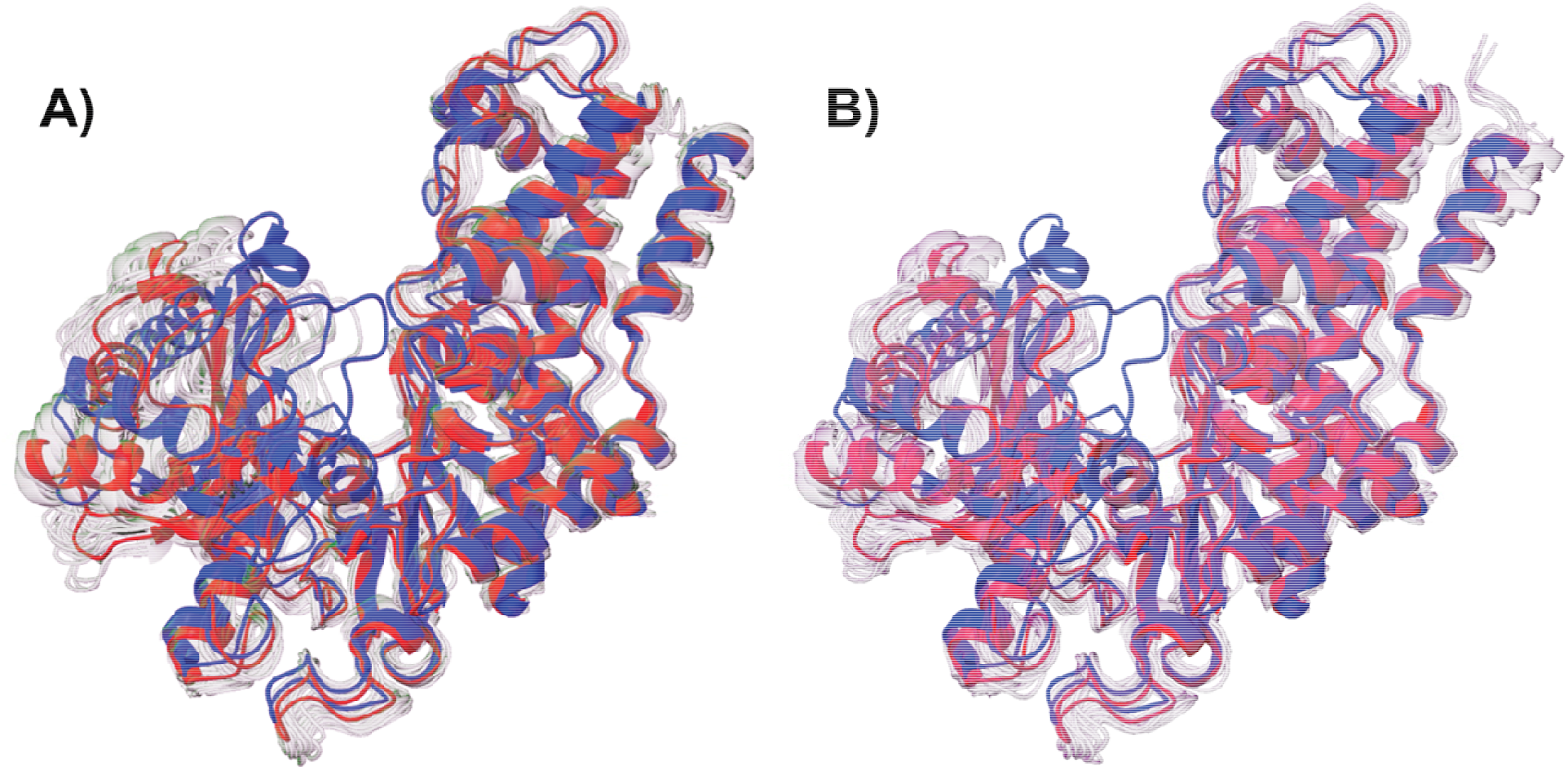
*Sc*Hxk2p SubPEx vs. brute-force simulations. The starting conformation (PDB 1IG8) is shown in red. The closed conformation (PDB 3O8M) is shown in blue. (A) SubPEx-sampled conformations identified using per-generation clustering. (B) Brute-force sampled conformations identified using traditional clustering.

## 5. Comparison with other methods

Aside from the WE approach, there are several other enhanced-sampling MD methods. The so-called “biased” methods encourage conformational transitions by altering the underlying free-energy landscape^27,78-86^. These methods are excellent for many use cases but are not ideal for our purposes. By altering the free-energy profile, some biased methods sample unphysical transition mechanisms/pathways^87-89^. In contrast, other simulation methods (including WE) take a directed, “unbiased” approach^90-96^. Though typically used to enhance whole-protein sampling, these methods can also sometimes identify novel pocket conformations^97,98^. Markov State Models (MSMs) are particularly appealing, but we note that WE captures both Markovian and non-Markovian dynamics^99,100^, tends to require less total simulation time^101^, and is not sensitive to the choice of lag time^99,100^.

SubPEx is fairly unique among applications of the WE methodology^33,99^. WE typically guides MD simulations towards a single target conformation, but SubPEx guides simulations towards increasingly dissimilar pockets. As there is no single optimally dissimilar pocket, the reaction coordinate does not have a single endpoint. Our approach is similar in some ways to the WExplore algorithm^100^, but focused on pocket shapes rather than protein conformations.

To our knowledge, only Johnson et al. have devised a method that prospectively focuses computational effort on sampling known pockets^102^. Though inventive, their approach drives the simulation towards the single largest pocket and so may fail to capture the full range of diverse pocket conformations. SubPEx enhances sampling along its entire reaction coordinate and so does not have this same limitation.

## 6. Conclusions

SubPEx uses weighted-ensemble (WE) path sampling^33^ to specifically enhance pocket sampling. Various experimental^103^ and computational^29,30,98,104^ methods can effectively identify both persistent and difficult-to-detect transient^105^ binding pockets. But these methods do not reveal the full range of ligand-accommodating conformations that such pockets adopt. Once a pocket has been identified, SubPEx allows researchers to better explore its conformational flexibility.

SubPEx is not well suited to all pockets. Some rigid pockets predominantly sample only one conformation and so are unlikely to benefit from enhanced sampling. Other pockets so readily interconvert between different conformational states that brute-force MD is sufficient to capture the full range of pocket flexibility. We expect SubPEx to be useful for studying pockets that only rarely transition between distinct pharmacologically relevant conformations.

In this work, we provided three examples that show how SubPEx can accelerate pocket sampling over brute-force MD. Of note, these SubPEx simulations did not start from crystal structures with small-molecule ligands bound in the respective orthosteric sites. Ligand binding frequently induces pocket conformational changes that cannot be easily predicted from ligand-unbound (*apo*) structures, yet such ligand-bound (*holo*) conformations are often most useful for drug discovery^106^. In other words, the very conformations that are most pharmacologically relevant can often only be observed if co-crystallized with an existing known ligand, complicating first-in-class discovery. Thankfully, per the population-shift model of binding^107-110^, *apo* simulations of sufficient length should capture *holo*-like conformations, even if only briefly. Indeed, we have leveraged *apo* simulations in several ensemble-based VS projects^16,17,19,21^. SubPEx accelerates the exploration of binding-pocket flexibility, expanding the utility of *apo* simulations as tools for generating pharmacologically useful conformational ensembles.

Though we here focus on *apo* pocket conformations, SubPEx may also be helpful in other contexts. For example, we can imagine other scenarios in which enhancing the sampling of specific protein regions (e.g., biologically important flexible loops) could be useful. Further, we have not yet explored whether SubPEx might be applied to simulations of ligand-occupied (*holo*) pockets; such simulations could reveal cryptic sub-pocket extensions of the primary *holo* pocket that may be exploited via lead optimization.

Finally, we note that in the present work, we ignore the weights that WESTPA rigorously tracks and reallocates to account for each walker’s contribution to the statistical ensemble^33,100,111^. This weighting scheme permits the calculation of thermodynamic and kinetic properties (e.g., transition rates)^33,89,112^, but it requires much longer, fully equilibrated WE simulations that have reached meaningful state probabilities. We prioritized accelerating conformational sampling over calculating transition kinetics between different conformational states. Users with different priorities can certainly apply the WESTPA framework’s useful analysis scripts to longer, equilibrated SubPEx simulations.

We are hopeful that SubPEx will help computational chemists efficiently incorporate protein flexibility into drug discovery (e.g., ensemble docking), even when transitions between pocket conformational states are rare. We release the software under the terms of the MIT license. A copy is available free of charge (without registration) from http://durrantlab.com/subpex/.

## 7. Supporting Information

Details regarding additional progress coordinates, simulations, and HSP90/NA/*Sc*Hexk2 analyses, including Table S1 and Figures S1-S6 (PDF).

## 8. Acknowledgments

We thank Kim F. Wong for help with debugging, as well as Lillian T. Chong, David R. Koes, Maria Kurnikova, Kevin Cassidy, Roshni Bhatt, Yuri Kochnev, and the Weighted Ensemble community for helpful discussions. This work was supported by the National Institute of Health (1R01GM132353-01A1); the University of Pittsburgh Center for Research Computing, RRID:SCR_022735 (supported by NSF OAC-2117681); and the NSF XSEDE program (bio200078). The content is solely the responsibility of the authors and does not necessarily represent the official views of the National Institutes of Health or the National Science Foundation. The funders had no role in the study design, data collection and analysis, decision to publish, or preparation of the manuscript.

