## supporting information for "Worth the weight: Sub-Pocket EXplorer (SubPEx), a weighted-ensemble method to enhance binding-pocket conformational sampling"

### 1. Additional Progress Coordinates

### 1.1 Jaccard distance

Aside from the progress coordinates described in the main text, we tested several additional coordinates. The first is based on the Jaccard Distance (JD) between two pocket volumes. Calculating JD leverages an approach similar to that used by the POVME algorithm^1,2^. The user provides the center and radius of the pocket to be sampled, which SubPEx uses to generate a sphere of equally spaced grid points. SubPEx then removes the points that come within 2.6 Å of any protein atom or that fall outside the convex hull defined by the protein Cα atoms. We chose a 2.6 Å cutoff because it is roughly equal to the van der Waals radii of water^3^ and hydrogen^4^, summed. Finally, SubPEx clusters the remaining points using DBScan, as implemented in the scikit-learn package (0.22.1)^5^, and retains only those points belonging to the largest cluster. This final field of points (FOP) occupies the binding pocket and defines its shape. To assess pocket dissimilarity, SubPEx then compares this FOP to the FOP of the reference (initial) pocket conformation,

$${JD}_{i}=1- \frac{\left| {FOP}_{i}\cap{FOP}_{ref} \right|}{\left| {FOP}_{i}\cup{FOP}_{ref} \right|}$$

where *JD_i_* is the Jaccard distance at walker *i*, *FOP_i_* is the field of points that occupy the walker binding pocket, *FOP_ref_* is the field of points that occupy the reference binding pocket, ${FOP}_{i}\cap{FOP}_{ref}$ is the set of points the two fields share in common, and ${FOP}_{i}\cup{FOP}_{ref}$ is the set of points in either or both fields. *JD_i_* thus ranges from 0 (pocket identity) to 1 (complete pocket dissimilarity).

### 1.2 Jaccard distance + pRMSD

Both the JD and pRMSD metrics are degenerate, meaning two distinct pockets can be equally dissimilar to the reference. To break this degeneracy, SubPEx also provides a two-dimensional progress coordinate involving both JD and pRMSD. While two distinct pockets may have similar JD metrics or similar pRMSD metrics, they are unlikely to be simultaneously similar in both metrics.

### 1.3 pRMSD + bbRMSD

The one-dimensional cRMSD progress coordinate described in the main text seeks to enhance both pocket and backbone flexibility, with a predominant focus on the pocket. As described below, we also considered a two-dimensional bbRMSD/pRMSD progress coordinate that encourages sampling of conformations that differ from the reference in terms of the backbone (bbRMSD) and pocket (pRMSD).

### 2. Technical details for additional simulations described in the Supporting Information

Table 1 provides the technical details for the brute-force and SubPEx simulations described in the main text. Table S1 provides the same information for the additional simulations described in the Supporting Information.

Table S1. Technical details for brute-force and SubPEx simulations. We used the minimal adaptive binning scheme^6^ for all SubPEx simulations except the HSP90 Amber20 bbRMSD/pRMSD simulation, which used a rectilinear binning scheme.

| Brute-force and SubPEx | | SubPEx Only | | |
| --- | --- | --- | --- | --- |
| System | MD Engine | Progress Coordinate | # Bins | Walkers/Bin |
| HSP90 (5J2V)^7^ | NAMD 2.14 | JD | 19 | 3 |
| HSP90 (5J2V)^7^ | Amber20 | bbRMSD/pRMSD | 1089 | 3 |
| HSP90 (5J2V)^7^ | NAMD 2.14 | bbRMSD/pRMSD | 108 | 3 |
| NA (2HTY:A)^8^ | Amber20 | JD/pRMSD | 68 | 5 |

### 3. HSP90: Additional Analyses

### 3.1 JD progress coordinates fails to adequately capture pocket flexibility

The main text describes how our HSP90 SubPEx simulation using the pRMSD progress coordinate failed to adequately capture pocket flexibility because it did not simultaneously enhance backbone flexibility. We also tested the one-dimensional JD progress coordinate, which similarly does not consider backbone flexibility. These efforts also failed to adequately capture pocket flexibility.

The JD SubPEx simulation started from the same equilibrated *apo* HSP90 (PDB 5J2V^7^) structure used in the pRMSD SubPEx simulations and ran for 50 generations (46.98 ns of cumulative simulation time and a maximal trajectory length of 1 ns). Figure S2A provides the JD progress-coordinate values per WE generation.

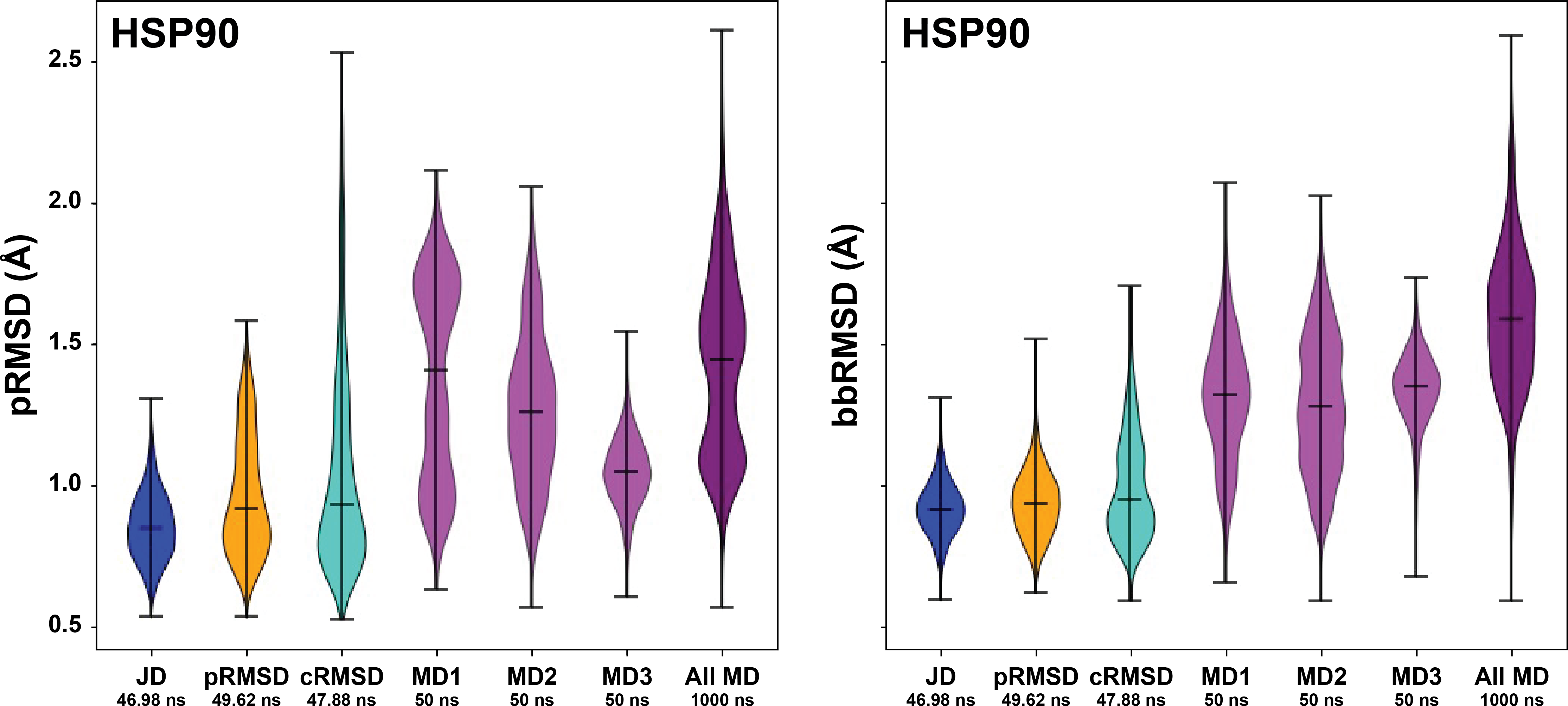

Figure S1. Violin plots comparing SubPEx HSP90 simulations with brute-force MD simulations. SubPEx simulations using the JD, pRMSD, and cRMSD progress coordinates are shown in blue, orange, and turquoise, respectively. In purple, three brute-force MD simulations (first 50 ns). In darker purple, all the frames of the brute-force simulations concatenated (1 μs total). Note that this figure is the same as Figure 1 in the main text, but with the JD-coordinate sampling included for comparison.

To compare the JD SubPEx simulations to brute-force HSP90 simulations, we considered the same three brute-force HSP90 simulations used to assess pRMSD SubPEx sampling (250 ns, 250 ns, and 500 ns, totaling 1 μs). We separately considered the first 50 ns of each brute-force simulation (to roughly match the cumulative simulation time of the SubPEx simulation) as well as the entire concatenated trajectory (1 µs) for reference. To assess the extent of pocket and backbone sampling, we calculated the pRMSD and bbRMSD of the simulated frames (auxiliary data).

Figure S1A shows that the JD SubPEx simulation failed to capture the pocket (and backbone) conformational sampling of the similar-duration brute-force simulations.

### 3.2 bbRMSD/pRMSD progress coordinate

In the main text, we describe our use of a one-dimensional progress coordinate (cRMSD) that considers both protein and backbone flexibility. We also explored whether a two-dimensional bbRMSD/pRMSD progress coordinate that similarly incorporates whole-protein backbone flexibility (bbRMSD) could also improve pocket sampling. This effort was less successful, so we discuss it only briefly here in the Supporting Information.

We ran a bbRMSD/pRMSD SubPEx simulation for 73 generations (cumulative simulation time of 253.74 ns, maximal trajectory length of 1.46 ns) and compared it to a similar brute force simulation. We assessed the extent of pocket and backbone sampling using the pRMSD and bbRMSD of the simulated frames as auxiliary data (independent of the progress coordinate). An analysis of the bbRMSD values (data not shown) suggested that bbRMSD/pRMSD SubPEx still sampled only limited backbone flexibility, even though bbRMSD contributes to the two-dimensional bbRMSD/pRMSD coordinate. Given this poor performance, we recommend using the 1D cRMSD progress coordinate instead of the 2D bbRMSD/pRMSD coordinate.

### 3.3 Progress-coordinate distributions for JD, pRMSD, and cRMSD HSP90 SubPEx runs over time

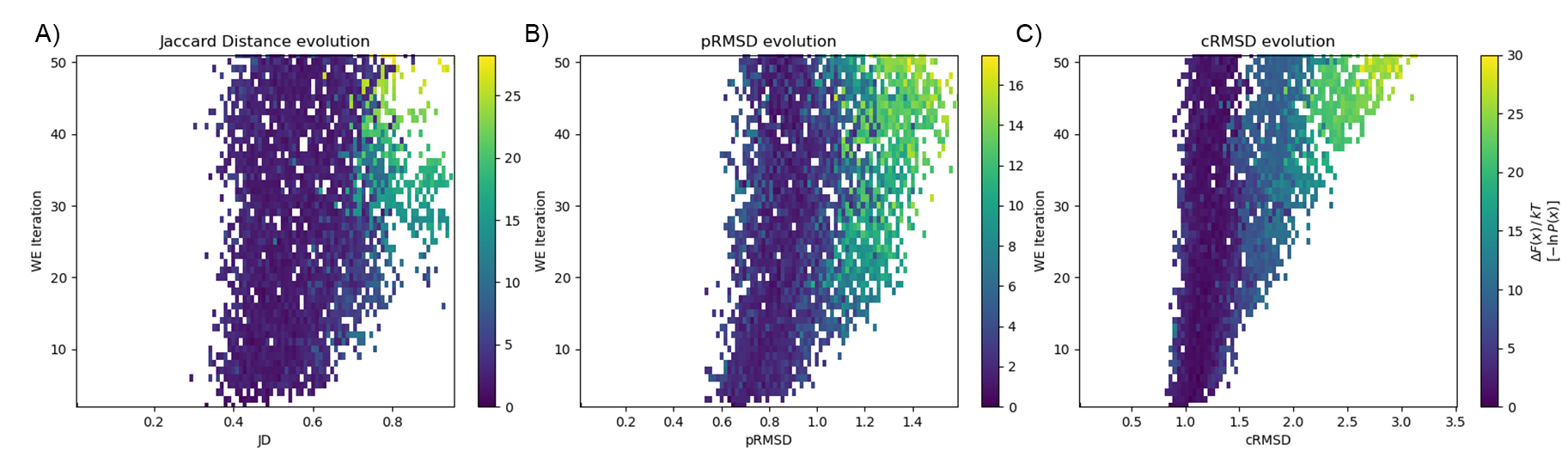

Figure S2. Probability distribution plots for three 50-generation SubPEx HSP90 simulations per WE generation (iteration). (A) JD progress coordinate. (B) pRMSD progress coordinate. (C) cRMSD progress coordinate. Note that the X axes and color schemes do not share the same scale.

We describe the pocket and backbone conformational sampling of the JD HSP90 SubPEx simulations above (Figure S1), and the pRMSD and cRMSD HSP90 SubPEx simulations in the main text (Figure 1). Figure S2 shows how each of these one-dimensional SubPEx runs sampled its respective progress coordinate per generation (iteration). The sampling of the JD SubPEx simulation stalled; only a few walkers reached higher JD values (i.e., approaching the maximum possible value of 1), with most of the probability between ~0.4 and ~0.65. In contrast, in the pRMSD SubPEx simulation, more walkers sampled higher progress-coordinate values, though progress along this coordinate was also limited (maximum value of only ~1.6 Å). In contrast, the cRMSD SubPEx simulations made sustained progress along its coordinate over the 50 generations shown, reaching a maximum value of ~3.1 Å.

### 3.4 SubPEx simulations compared to crystallographic references (per RMSD)

To supplement the principal component analysis (PCA) presented in the main text, we further evaluated SubPEx and brute-force simulations by calculating the pocket heavy-atom RMSDs between each simulation frame and the crystallographic reference structures 4EFT^58^ and 4YKR. These structures capture the pocket-adjacent D102-G114 residues in helical and loop conformations, respectively. We visualized these RMSD values using a 2D histogram (Figure S3).

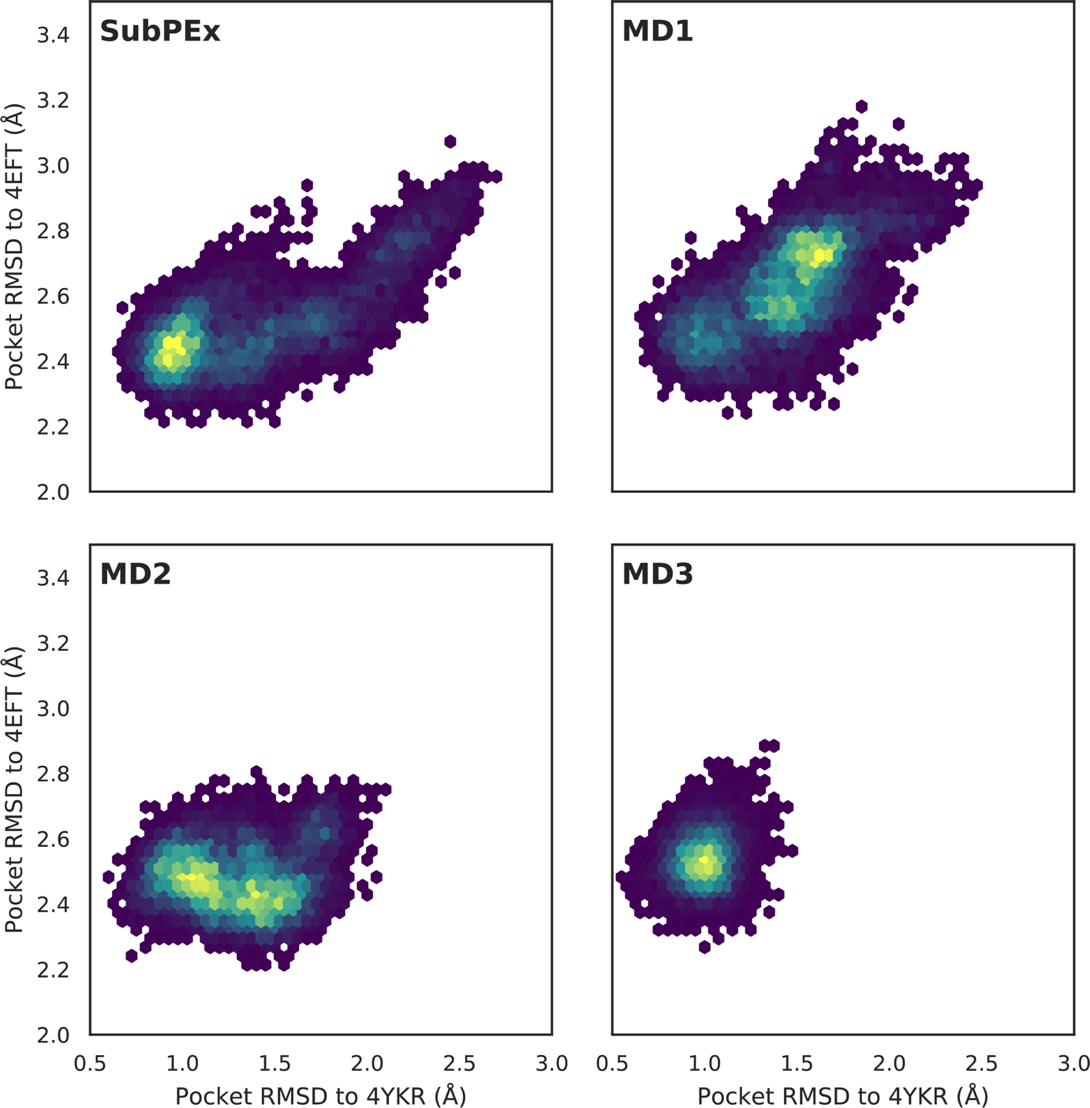

Figure S3. Comparing HSP90 simulation frames and crystallographic references (RMSD). For each (SubPEx + three brute-force) simulation, we calculated the heavy-atom pocket RMSD between all pruned frames and two crystallographic reference structures: 4YKR (x-axis), with D102-G114 in a loop conformation; and 4EFT (y-axis), with D102-G114 in a helical conformation. These values were binned into 2D histogram for easy visualization. All four panels capture 102.6 ns of cumulative simulation time.

One metric of success is how well a given technique samples known pocket conformations. Here the SubPEx and brute-force simulations performed comparably. Interestingly, none of the simulations perfectly captured the helical conformation typical of the 4EFT structure, although all simulations came within 2.3 Å of that conformation at some point. The simulations were even more successful at sampling the 4YKR loop conformation (within 0.7 Å).

A second metric of success is how well a given technique samples pocket conformations that are notably different from any known conformation, potentially revealing novel pocket shapes that can accommodate unique ligands. Here SubPEx performs particularly well. It samples a broader range of conformations than the second (MD2) and third (MD3) brute-force simulations. Although the first brute-force simulation (MD1) samples a similarly sized region within this RMSD space, it predominantly samples conformations that differ substantially from the two known crystallographic references (e.g., the distribution peak is far from the origin). In contrast, SubPEx primarily samples conformations near known crystal structures, while also exploring regions distant from these references. SubPEx thus arguably better reveals pocket diversity while staying true to experiment.

### 4. NA: Additional analyses

### 4.1 Limitations of JD/pRMSD progress coordinate

To assess how well SubPEx improves sampling of the flexible NA 150-cavity, we also performed SubPEx simulations of the open-150-cavity structure (PDB 2HTY:A^8^) using a two-dimensional JD/pRMSD progress coordinate (Table S1) for 75 total generations. The maximal SubPEx trajectory length (i.e., the longest continuous simulation time of any single walker from the first to last generation) was 1.50 ns. The aggregate SubPEx simulation time (i.e., the total time sampled by all walkers across all generations) was 360.3 ns.

The SubPEx simulation samples more of the progress-coordinate space than the brute-force simulations, especially in the lower pRMSD and JD regions (Figure S4). However, the brute-force MD simulations sample more of the higher pRMSD and JD values. It is also noteworthy that the brute-force MD simulations sample around what seems to be an energy minimum, but the SubPEx simulation is not stuck in the minimum.

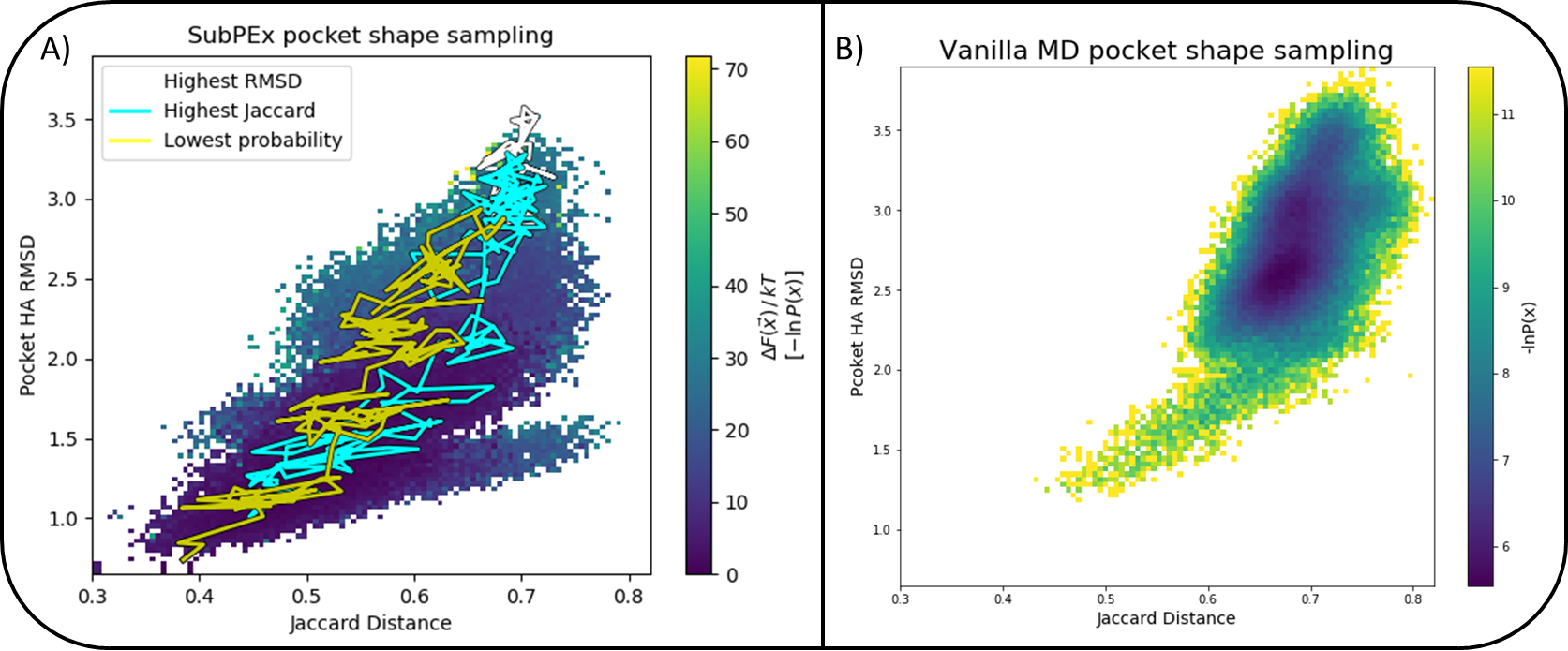

Figure S4. NA JD/pRMSD SubPEx sampling compared to brute-force sampling. (A) Two-dimensional probability distribution as a function of the JD (x-axis) and pRMSD (y-axis). In a yellow trace, we show the path the lowest probability walker took; in blue and white, the walker with the highest JD and pRMSD, respectively. The cumulative simulation time is 360.3 ns. (B) Two-dimensional probability distribution as a function of the JD (x-axis) and pRMSD (y-axis) of the brute-force MD simulations (total simulation time 1 μs), with the probability shown as counts (colors are inverted compared to A).

To determine why the SubPEx simulations do not adequately sample higher pRMSD conformations in this case, we compared the pRMSD and bbRMSD values of the SubPEx- and brute-force sampled frames (Figure S5). pRMSD and bbRMSD are useful for assessing the extent of pocket and whole-protein sampling, respectively. Although SubPEx does not sample conformations with pRMSD values as high as those of the brute-force simulations, SubPEx does more thoroughly sample pockets with lower pRMSD values (e.g., the distribution of pRMSD values is more even; Figure S5, on the left). However, the SubPEx simulations have limited backbone sampling relative to the brute-force simulations (Figure S5, on the right).

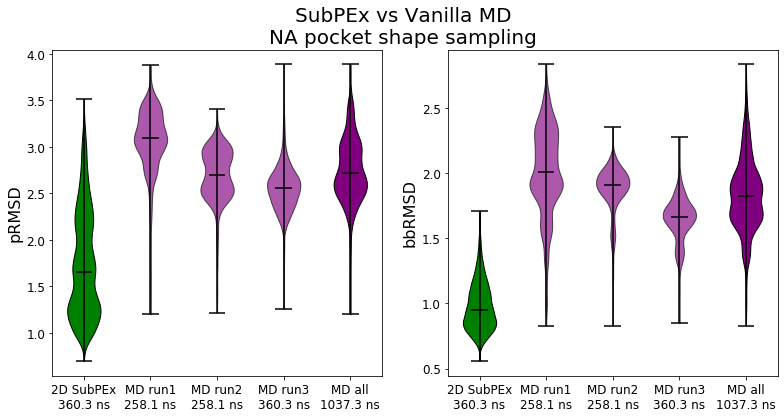

Figure S5. Violin plots of the pRMSD and bbRMSD distributions for NA JD/pRMSD (2D) SubPEx and brute-force simulations. In green, the JD/pRMSD SubPEx simulations (360.3 ns cumulative simulation time). In purple, the brute-force MD simulations. In dark purple, the three MD simulations, concatenated. Only the first 360.3 ns of the third brute-force MD simulation are included in this analysis to enable proper comparaison with the SubPEx simulation. The other two MD simulations are presented without truncation since they are shorter than 360.3 ns.

### 4.2 Brute-force and SubPEx NA simulations starting from the closed conformation

In the main text and above, we describe NA SubPEx simulations that start from an open 150-cavity NA structure (PDB 2HTY^8^). We also performed similar simulations that started from a closed 150-cavity NA structure (PDB 2HU4:A^8^). Unfortunately, two of the three systems only equilibrated after ~50 ns. Consequently, the SubPEx simulations did not reach the same high pRMSD values as the brute-force simulations because SubPEx used its enhanced sampling to equilibrate the closed system. We mention this outcome here for completeness’ sake and as a cautionary tale. Users should take care to ensure that their systems are fully equilibrated before beginning SubPEx runs.

### 5. *Sc*Hxk2: Additional analyses

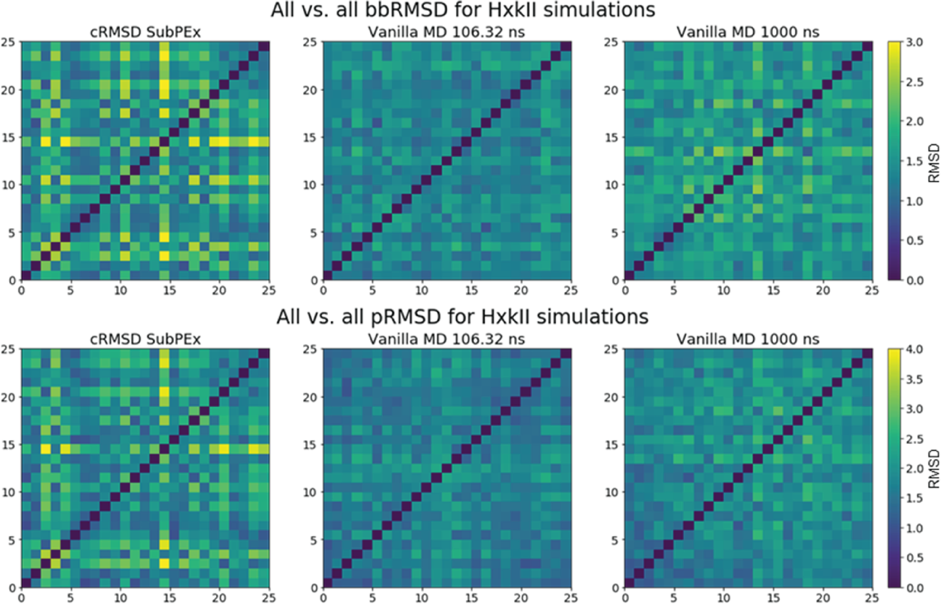

Figure S6. All vs. all RMSD of clustered protein conformations for the brute-force and cRMSD SubPEx *Sc*Hxk2 simulations.
